# Single-molecule simultaneous profiling of DNA methylation and DNA-protein interactions with Nanopore-DamID

**DOI:** 10.1101/2021.08.09.455753

**Authors:** Seth W. Cheetham, Yohaann M. A. Jafrani, Stacey B. Andersen, Natasha Jansz, Michaela Kindlova, Adam D. Ewing, Geoffrey J. Faulkner

**Affiliations:** Mater Research Institute – University of Queensland, TRI Building, Woolloongabba QLD 4102, Australia; Australian Institute for Bioengineering and Nanotechnology, University of Queensland, St Lucia, Australia; Genome Innovation Hub, The University of Queensland, Brisbane, QLD 4072, Australia; Queensland Brain Institute, University of Queensland, Brisbane QLD 4072, Australia

## Abstract

We present Nanopore-DamID, a method to simultaneously detect cytosine methylation and DNA-protein interactions from single molecules, via selective sequencing of adenine-labelled DNA. Assaying LaminB1 and CTCF binding with Nanopore-DamID, we identify escape from LAD-associated repression of hypomethylated promoters amidst generalised hypermethylation of LaminB1-associated regulatory elements. We detect novel CTCF binding sites in highly repetitive regions, and allele-specific CTCF binding to imprinted genes and the active X chromosome. Nanopore-DamID highlights the importance of DNA methylation to transcription factor activity.

## Main

Cytosine methylation modulates the interactions of chromatin-associated proteins with DNA^1^. DNA adenine methylase identification (DamID) is an approach to profile DNA-protein interactions^2,3^. In DamID, the *E. coli* adenine methylase Dam is fused to a chromatin-associated protein (**Fig. 1a**). The Dam-fusion protein methylates adenines in GATC motifs proximal to the site of chromatin-association. Methylated GATC sites are then cleaved by the methylation-specific restriction enzyme DpnI, followed by adaptor ligation, amplification and Illumina sequencing^4^. DamID has been adapted in various model organisms^2,5–9^ to detect chromatin accessibility^10^, RNA-chromatin interactions^11^, chromatin topology^12,13^, chromatin state^14,15^ and nuclear lamina interactions^16^. Despite the utility of DamID, the approach has drawbacks. Each molecule must have GATC methylation on both ends to be amplified, reducing resolution and rendering DamID incompatible with bisulfite sequencing.

**Figure 1:**
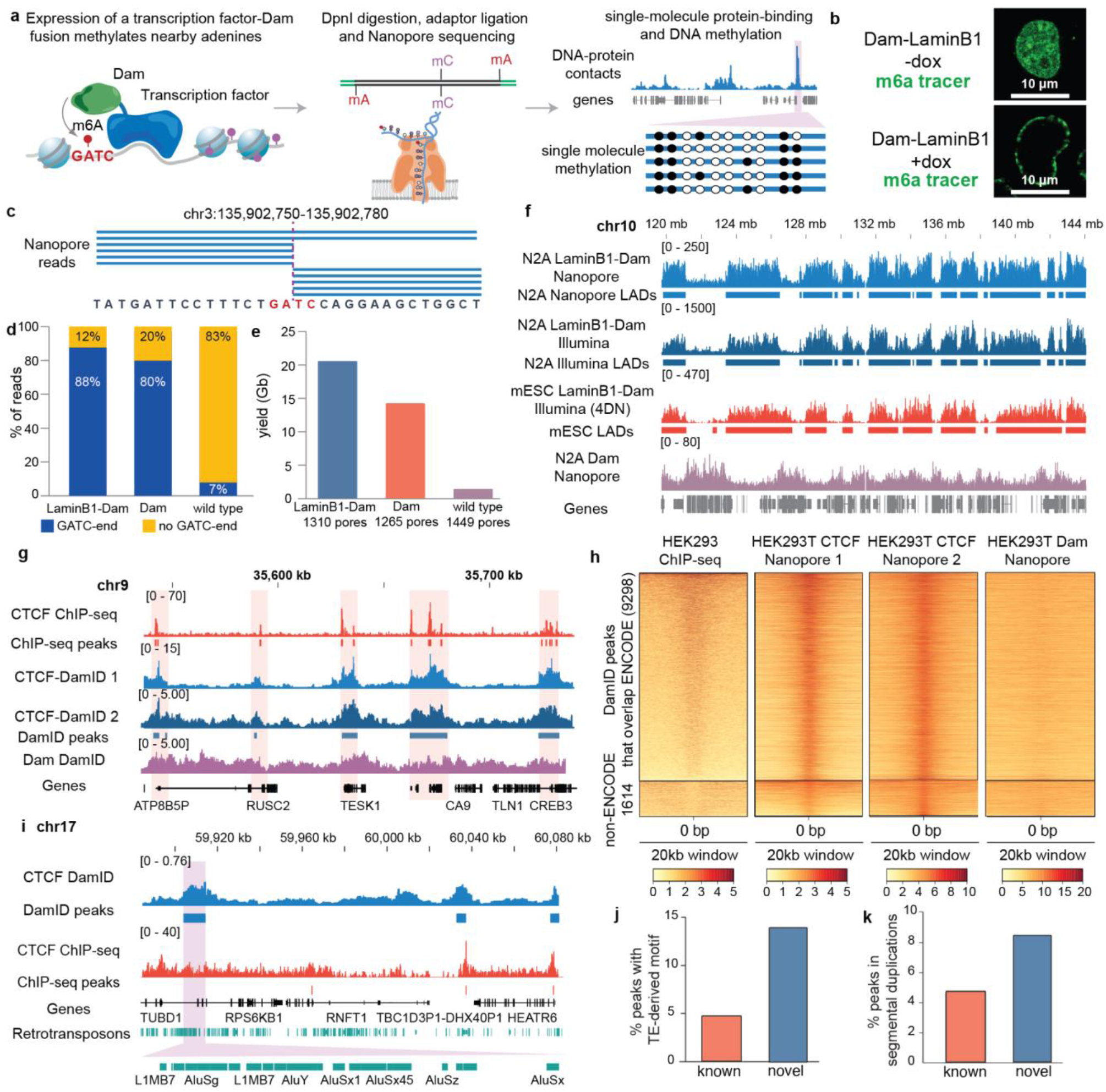
Nanopore-DamID identifies Lamin-associated domains and CTCF binding sites. **a**, In Nanopore-DamID, a fusion Dam-chromatin-associated protein is expressed in cells and methylates adenines in GATC motifs proximal to sites of DNA-protein interaction. Genomic DNA is isolated from cells, dephosphorylated and digested with DpnI. Digested DNA ends are A-tailed and then Nanopore sequencing adaptors are ligated. Nanopore sequencing identifies DNA-protein contacts and CpG methylation on single DNA molecules. **b**, Upon doxycycline treatment, the m6A-tracer (an inactive DpnI fragment fused to GFP) localises to the nuclear lamina in cells transfected with TetON-Dam-LaminB1. **c**, Example of Nanopore-DamID reads that terminate at a GATC motif. **d**, Most reads from Nanopore-DamID of Dam-LaminB1 and Dam expressing cells, but not wild type cells, terminate at a GATC motif. **e**, The yield of sequencing from wild type cells is reduced compared to Dam expressing cells. **f**, Nanopore-DamID of LaminB1-Dam-expressing N2A cells shows enrichment in large lamin-associated domains similar to Illumina DamID and published mESC Illumina-DamID^26^. In contrast, untethered Dam occupancy is enriched in inter-LAD regions. **g**, Profiles of CTCF occupancy from ChIP-seq (ENCODE) and Nanopore-DamID **h**, Enrichment of ChIP-seq signal, but not untethered Dam over most DamID peaks. **i**, Example of a CTCF-binding site in a transposable element (TE)-rich region not detected by ENCODE. **j**, Enrichment of TE-derived CTCF motifs in peaks not detected by ENCODE **k**, Enrichment of peaks not detected by ENCODE in segmental duplications.

Oxford Nanopore Technologies (ONT) long-read sequencing can directly detect both cytosine and adenine methylation, including in regions refractory to short-read analysis^17–19^. Exogenous promiscuous adenine methylases have recently been used to profile chromatin accessibility^20^ and, by fusion to protein-A, regions of transcription factor occupancy bound by a cognate antibody^21,22^. However, direct detection of adenine methylation requires high genome-wide coverage, which can be prohibitively expensive in mammals, and most eukaryotes. Here, using LaminB1 and CTCF as examples, we develop a Nanopore-based DamID which enables rapid, cost-effective single-molecule profiling of transcription factor-cytosine methylation interactions.

Nanopore sequencing can be used to selectively analyse adaptor-ligated DNA without purification or amplification, for example to enrich fragments cleaved by Cas9^23^. Thus, if adaptors are only ligated to DNA cleaved by the methylated-GATC-specific enzyme DpnI, these fragments can be selectively sequenced from a heterogeneous mixture of largely unligated fragments (**Fig. 1a**). To enable selective sequencing of Dam-methylated fragments, we stably transduced the mouse neuroblastoma cell line, N2A, with a doxycycline-inducible Dam-LaminB1 fusion protein or untethered Dam expression cassette (**Supplementary figure 1a-b)** and constitutively expressed m6A-tracer^24^ (GFP fused to catalytically inactive DpnI, **Supplementary figure 1c**). Characteristic fluorescent rings at nuclear lamina were evident in cells upon doxycycline treatment, confirming that Dam-LaminB1 can efficiently modify GATC sites in lamin-associated chromatin (**Fig. 1b**).

To simultaneously detect DNA-protein interactions and CpG methylation on single molecules, we expressed Dam-LaminB1, or untethered Dam, in N2A cells and extracted genomic DNA. After de-phosphorylation, methylated GATC fragments were cleaved with DpnI and Nanopore adapters ligated (see **Supplementary figure 2** for size distribution). Library preparation here takes <1.5 days. From sequencing of LaminB1 and Dam-alone libraries on single MinION flow cells, we generated 10.8 million and 3.77 million reads respectively, which as expected terminated primarily at GATC motifs (**Fig. 1c-d**). 89% of LaminB1 and 80% of untethered Dam reads terminated with a GATC motif (**Fig. 1d**) and were thus derived from a DpnI-cleaved Dam-methylated site (**Table S1**). Indeed, *de novo* motif enrichment analysis on read termini identified profound enrichment of GATC motifs (p=3.3e-703, **Supplementary figure 3**). In contrast, only 7% of reads from wild type N2A cells subjected to Nanopore-DamID terminate at GATC motifs (**Fig. 1d**). Furthermore the yield of bases from wild type cells was greatly reduced (<10×) compared to Dam-expressing cells, reflecting minimal endogenous adenine methylation or minor cleavage of unmethylated GATC sites (**Fig. 1e**).

Nanopore reads were enriched in the broad lamin-associated domains (LADs) characteristic of LaminB1 occupying 55% of the mouse genome^16,25^ (**Fig. 1f**) whilst untethered Dam localised to open chromatin between LADs (inter-LADs or iLADs, **Fig. 1f**) and was negatively correlated with LaminB1 occupancy (inter-LADs or iLADs, Spearman correlation=-0.25). To compare Nanopore and Illumina-based approaches we sequenced conventional LaminB1-DamID libraries from N2A cells. LaminB1-occupancy profiles were comparable between technologies and similar to those obtained previously from mouse embryonic stem cells (mESCs)^26^ (**Fig. 1f**). LaminB1 profiles clustered by cell-type rather than sequencing technology (**Supplementary figure 4)**. Notably, the higher median read lengths of Nanopore-DamID (>1000bp, **Table S1, Supplementary figure 2**) enabled identification of DNA-protein contacts (4.23-8.27× more bases) in regions that were unmappable for short (75bp) reads (for an example see **Supplementary figure 5**). LADs were identified with a previously developed hidden markov model by comparing LaminB1-Dam signal to Dam-alone^27^. Nanopore-DamID LADs overlapped extensively with Illumina-DamID LADs (Jaccard metric=0.89, p<0.001 Genome Association Test^28^) and LADs identified from a published mESC LaminB1 DamID dataset^26^ (Jaccard metric=0.75, p<0.001 Genome Association Test), although some LADs were cell-type-specific (**Supplementary figure 6**). These results confirm that selective sequencing of adenine methylated DNA, without amplification or purification, is methodologically simple, accurate, and cheaper than approaches where whole genome sequencing is required^22^.

To determine if Nanopore-DamID can selectively sequence adenine-methylated DNA from a heterogeneous mixture of transgenic and wild type cells, we mixed 10% LaminB1-Dam expressing cells (1×10^5^ cells) with 90% wild type cells and performed Nanopore-DamID. We found the resulting profiles from diluted cells were highly comparable to those generated from undiluted cells (**Supplementary figure 7a-b**, Spearman correlation=0.85). Identified LADs were highly similar to those from undiluted cells (Jaccard metric=0.95) demonstrating that Nanopore-DamID can be used to profile a selected cell population in a mixture of unlabelled cells, and on as few as ∼1×10^5^ labelled cells.

To determine if Nanopore-DamID can detect transcription factor DNA binding, we performed Nanopore-DamID on the methylation-sensitive^29^ transcription factor CTCF in HEK293T cells. For this assay, we use a non-promiscuous Dam mutant, DamN126A^30^. As in previous experiments, >80% of reads terminated at GATC sites (**Table S1**) and Nanopore-DamID CTCF profiles agreed strongly between replicates and with known CTCF binding sites (**Fig. 1g-h**). We did not detect an enrichment of untethered DamN126A at CTCF sites, in contrast to previous studies using wild type Dam^31^, likely due to the reduced affinity of DamN126A for accessible chromatin^30^. Of the 10,912 CTCF binding sites identified, 9,298 (85%) overlapped with ENCODE CTCF sites (from all ENCODE cell lines, N=231,761, Genome Association Test p<0.001, 3.49-fold enrichment) and 5,746 (52.7%) overlapped with HEK293 CTCF ChIP-seq peaks (N=38,394 ENCODE, Genome Association Test p<0.001, 6.3-fold enrichment). Whilst some of the peaks not previously detected by ENCODE may have been due to inherent differences between DamID and ChIP-seq^3,8^ others appeared due to the longer read length of Nanopore-DamID (>1000bp, **Table S1**) resolving binding sites in repetitive regions^32^. Indeed, peaks not detected by ENCODE were enriched in motifs derived from transposable elements (**Fig. 1i-j)**. Similarly, peaks only detected by Nanopore-DamID were enriched in segmentally duplicated regions (**Fig. 1k**), and particularly in large segmental duplications on chromosome one (80/136 novel peaks). These duplications contained, for instance, NOTCH2 and SRGAP2 copies regulating cortical development^33–36^ (**Supplementary figure 8**). Thus, Nanopore-DamID can resolve cryptic transcription-factor binding sites in repetitive regions refractory to shorter read lengths.

To study interactions between LaminB1 and DNA methylation, we identified CpG methylation from the LaminB1-Nanopore-DamID reads. Consistent with previous reports^38,39^ we found that, despite the heterochromatic structure of LADs, CpG methylation was globally reduced in these regions compared to non-Lamin-associated DNA, (inter-LADs, iLADs) (**Fig. 2a-b, Supplementary figure 9**). However, CpG islands and, most strikingly, the regions surrounding (± 150bp) transcriptional start sites were resistant to LAD hypomethylation (**Fig. 2a-b**). The maintenance of high methylation at LAD TSSs suggests that, despite tethering to the nuclear periphery, DNA methylation at the proximal promoter region may prevent aberrant heterochromatic gene expression in LADs. To determine the relationship between Lamin-interactions, DNA methylation and gene expression, we analysed previously published RNA-seq from N2A cells^40^. As expected, genes in LADs were almost completely silenced, with the exception of a subset of genes with hypomethylated TSSs (aggregate methylation <0.3, **Fig. 2c-d**) that have significantly higher expression (p=2.27e-22, Student’s t-test compared to hypermethylated LAD TSSs), escaping LAD-associated repression. Silent (counts per million, CPM <10) gene TSSs in LADs were highly methylated, as were silent iLAD genes (**Fig. 2e**). While almost all robustly expressed (CPM >10) LAD genes were profoundly hypomethylated, not all hypomethylated genes were expressed (**Fig. 2e**), suggesting that hypomethylation is necessary but not sufficient for escape from Lamin-associated repression. Crucially, the single-molecule nature of Nanopore-DamID clearly resolves hypomethylated Lamin-associated alleles, as distinct to hypomethylated alleles present in cells where these loci are not Lamin-associated.

**Figure 2:**
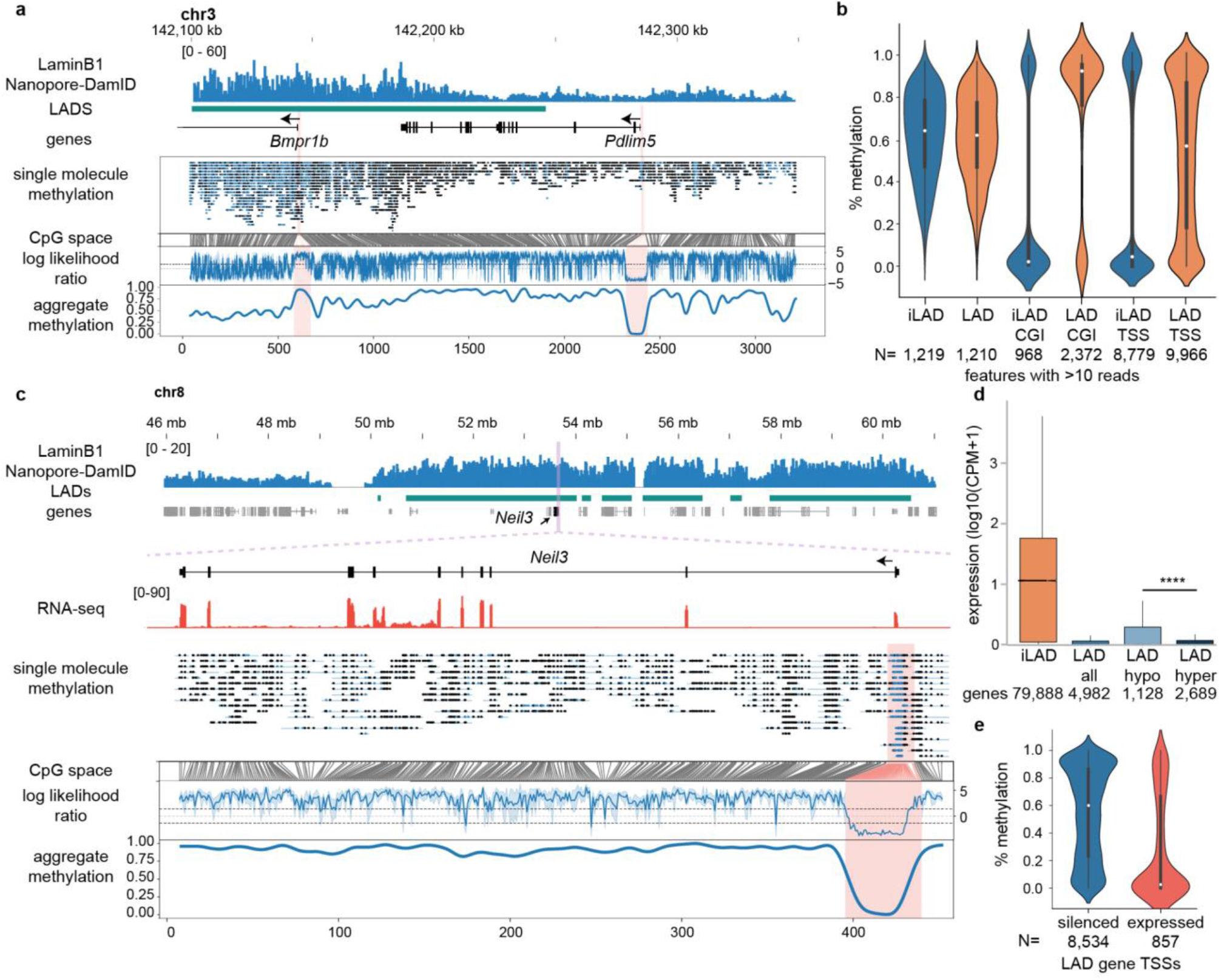
Nanopore-DamID identifies escape from repression by CpG hypomethylated genes in lamin-associated DNA. **a**, CpG-methylation from Lamin B1 Nanopore-DamID data visualised with Methylartist^37^ reveals a hypomethylated LAD on chromosome 3 containing hypermethylated LAD promoters. **b**, Genome-wide analysis of methylation within LADs or between LADs (iLADs) reveals hypomethylation of LADs, other than at CpG islands and TSSs (300bp window), which maintain higher methylation. **c**, A hypomethylated expressed gene in a LAD on chromosome 8. **d**, Comparison of expression of genes within LADs. Hypomethylated TSS (<30% methylation) were more highly expressed than hypermethylated genes (>70% methylation),p=2.27e-22, Student’s t-test. **e**, Genes expressed (>10 CPM) from LADs were almost universally hypomethylated.

We next used Nanopore-DamID to determine whether CTCF binding to CpG-containing CTCF motifs was methylation sensitive (**Fig. 3**). CTCF-bound motifs were almost universally demethylated (median methylation=0, N=53), whilst unbound motifs were largely methylated (median methylation=0.75), consistent with binding motif methylation being a key determinant of CTCF-binding^29^ (**Fig. 3a**). Notably, a minority of CpG-containing CTCF motifs were bound even in the presence of motif methylation (14%) suggesting a rare methylation-independent mode of binding. We previously generated Nanopore whole genome-sequencing (WGS, ∼28×) from HEK293T cells^41^, and here we also performed Illumina WGS to phase the nanopore reads. We identified 5,047 differentially methylated regions (DMRs) that displayed allele-specific methylation, 57% of which localised to the X chromosome due to differential methylation of the active and inactive X chromosomes (**Fig. 3b**). CTCF-Nanopore-DamID reads were significantly less methylated than unenriched whole genome-sequencing at DMRs (autosomal DMRs p=0.029, X chromosome DMRs, p=0.00049, Student’s t-test) reflecting increased CTCF interactions on the hypomethylated alleles (**Fig. 3c**). At the XIST locus, which controls X chromosome inactivation the phased WGS reads identified a demethylated and methylated allele corresponding to the active and inactive X chromosome. We found that all CTCF Nanopore-DamID reads at the XIST promoter were demethylated demonstrating that CTCF binding occurred exclusively at the active hypomethylated allele (**Fig. 3d**). Similarly, at the autosomal maternally imprinted locus KCNQ1OT1, CTCF-bound alleles were largely demethylated (**Fig. 3e**), demonstrating the capacity of Nanopore-DamID to detect single-molecule, allele-specific transcription factor-DNA methylation interactions.

**Figure 3:**
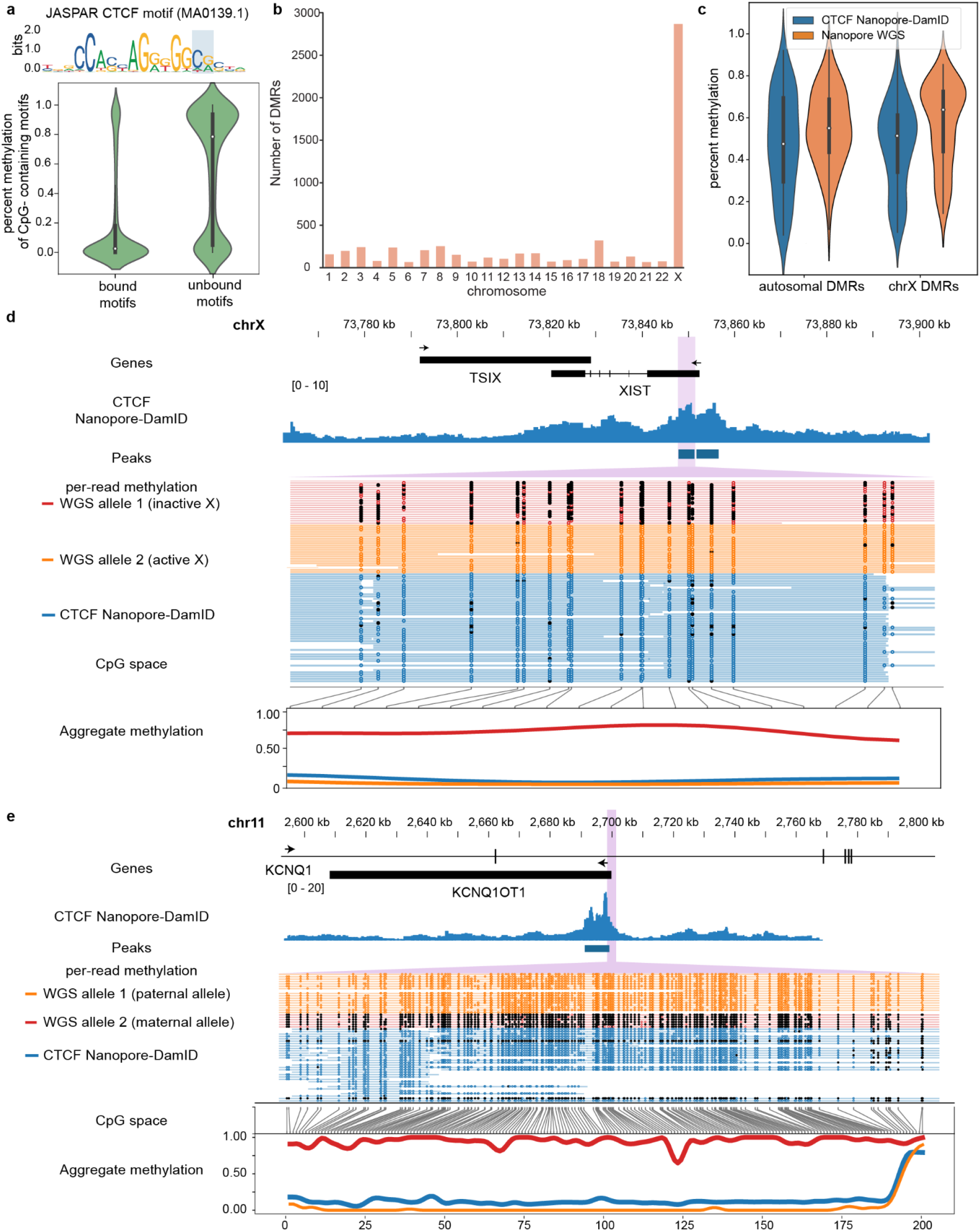
Nanopore-DamID identifies allele-specific transcription factor interactions. **a**, Methylation of CTCF motifs containing a CG dinucleotide in Nanopore-DamID peaks (bound) compared to non-peak (unbound) motifs. **b**, Enrichment of DMRs identified from HEK293T Nanopore WGS on the X chromosome. **c**, Comparison of the methylation of reads from CTCF Nanopore-DamID and Nanopore WGS at DMRs. **d**, Example of an allele-specific CTCF-binding site at the X-inactive specific transcript (XIST) locus. All Nanopore-DamID reads were demethylated and thus derived from the inactive X allele. **e**, Example of an allele-specific binding CTCF-binding site at the maternal imprinted gene KCNQ1OT1. Almost all Nanopore-DamID reads were demethylated and thus derived from the demethylated paternal allele.

Overall, Nanopore-DamID selectively sequences Dam-labelled native DNA, enabling co-detection of DNA-protein contacts and DNA methylation. This approach could be combined with other adenine methylase-based approaches^20–22^ to increase sequencing depth over methylated regions. Compared to existing approaches, Nanopore-DamID is less labour-intensive, enriches labelled regions, enables coverage of regions with low mappability, and enables detection of allele-specific interactions. Here, Nanopore-DamID identified escape of hypomethylated genes from Lamin-associated repression, as well as allele-specific CTCF interactions on X chromosome and at imprinted loci. Nanopore-DamID could in the future be employed to elucidate interactions between CpG methylation, or other DNA base modifications, with chromatin-associated proteins that regulate eukaryotic gene expression.

## Methods

### Transgenic cell line preparation

1×10^5^ N2A or HEK293T (ATCC) cells were seeded in one well of a 6-well plate in 2mL of DMEM complete media (Gibco) supplemented with 10% Fetal Bovine Serum, 1% PenStrep and 1% L-glutamine, and maintained at 37°C with 5% CO_2_ in a humidified incubator. 0.2μg Super piggyBac Transposase Expression Vector, 0.25μg m6A-tracer and 0.25μg XLONE-Dam constructs (BlastR) were diluted in 100μl of Opti-MEM (Thermo Fisher). 2.1μl of FuGENE-HD® (Promega) was added and the reaction was briefly vortexed and incubated at room temperature for 10min prior to addition to the cells. The transfection mix was added dropwise to the well and incubated for 24hr. After 24hr the media was changed and the cells were selected for seven days in media containing 2.5μg/ml blasticidin.

### Imaging

1×10^5^ transgenic N2A cells were seeded in two 35mm glass bottom dishes (Cellvis™). Dam-LaminB1 fusion protein expression was induced with 2μg/ml doxycycline in one of the dishes. The m6A tracer in cells of both dishes were visualised and imaged at 488nm on an Olympus FV3000 microscope at 120X magnification with a 60X oil immersion objective followed by a 2X digital zoom. Images were analysed via Fiji^42^.

### Nanopore DamID

Dam fusion protein expression was induced with 2μg/mL doxycycline and genomic DNA (gDNA) was extracted from one million cells cultured in Nunc™ T75 flasks using an NEB Monarch® gDNA extraction kit according to manufacturer’s instructions, with the exception of using inversion rather than vortexing for mixing. 2.5μg of gDNA was dephosphorylated by the addition of 5μl of 10X NEB Cutsmart® buffer and 1μl of QuickCIP (NEB) in a total volume of 48μL. The reaction was mixed by gentle flicking and incubated at 37°C for 30 minutes. The QuickCIP was then heat inactivated at 80°C for 5 min. 2μl of DpnI (NEB) was added to the reaction, mixed by gently flicking and incubated at 37°C overnight. DNA was A-tailed by addition of 1μL 10mM dATPs and 1μL of Klenow exo- (NEB) and incubation at 37°C for 1hr. The digested DNA was purified using the Qiagen QiaQuick™ PCR Purification kit (which purifies fragments <10kb) and eluted in 60μL of H_2_O. The DNA was prepared for ONT sequencing with the DNA Ligation Sequencing Kit (LSK110), starting from the adaptor ligation step and using the Short Fragment Buffer.

### Illumina DamID

Illumina-DamID-seq was performed as previously described with minor modifications^4^. Briefly, gDNA was extracted using the NEB Monarch gDNA extraction kit and digested overnight with DpnI (NEB). DamID adaptors were then ligated to DpnI-digested DNA. The adaptor-ligated DNA was digested with DpnII (NEB) to cleave fragments containing internal unmethylated GATC sites. Dam-methylated DNA was then amplified by PCR with MyTaq polymerase (Bioline®), sonicated to an average size of 300bp and the DamID adaptors were removed by AlwI (NEB) digestion. 1µg of sonicated DNA was prepared for sequencing using the NEBNext® Ultra™ II DNA Library Prep Kit for Illumina® without size selection and amplified with 3 cycles of PCR. Libraries were sequenced on the Illumina® NextSeq platform using single-end 75bp chemistry.

### DamID analysis

Nanopore reads were aligned to the mouse genome (mm10) using Minimap2^43^ with the options -a -- MD --cs=long -x map-ont. Mapped reads were filtered for map quality using SAMtools^44^ view -q 10 and converted to bed format using bedtools bamtobed and bigwig format using deepTools^45^ bamCoverage. The coordinates of the ends of reads were extracted using awk, extended to 100bp and converted to FASTA using bedtools getfasta. The sequences surrounding read ends were surveyed for enriched 4bp motifs using MEME-ChIP^46^. GATC motifs in the mouse genome (mm10) were extracted using the damidseq_pipeline^47^. The intersection of Nanopore read ends with GATC motifs was calculated using bedtools intersect. Short-read DamID was mapped using minimap2 ^43^ and filtered for alignment quality using samtools view -q 10. CpG methylation was called from Nanopore-DamID data using megalodon v2.2.9 with the model dna_r9.4.1_450bps_modbases_5mc_hac_prom.cfg. DNA methylation was visualised using Methylartist^37^ segplot and locus.

### Lamin-associated domains

Lamin-associated domains were identified by binning reads from LaminB1-DamID samples into 20 kb bins using bedtools coverage. Ratios of LaminB1-Dam/(Dam+1) were calculated using awk and Lamin-associated domains called using HMMt^27^. Consensus LADs (between the Nanopore-DamID undiluted and 1:1/10 dilution and between the two Illumina replicates) were determined using intersectBed.

### RNA-seq analysis

N2A wild type RNA-seq reads^40^ were downloaded (GSE140357) and mapped to the mouse genome (mm10) using STAR^48^ using default settings. Refseq transcripts were quantified using featurecounts^49^ with the options -t “exon” -O -Q 10.

### Motif analysis

CTCF motifs in the human genome (hg38, MA0139.1) were downloaded from JASPAR^50^. Motifs containing CpG dinucleotides were identified by conversion to fasta using bedtools getfasta and grep. Peaks containing only motifs with CpGs were identified by intersectBed. A random control set of motifs was identified by shuffling non-peak CpG motifs. Motifs within transposable elements (L1, SVA, *Alu* and HERV families) were identified by bedtools intersect against a subset of the UCSC Repeatmasker track^51^.

### Variant identification

HEK293T cells (p5) were grown to 70-85% confluency in the T75 flask, then washed with PBS and lifted with trypsin. Pelleted cells were lysed and genomic DNA was isolated using phenol-chloroform extraction protocol. DNA was quantified using a Quibit and prepared for sequencing using the Truseq(R) Nano kit at Macrogen. Sequenced on a HiSeq X using 150bp paired-end chemistry. Reads were aligned to hg38 using bwa-mem2^52^ version 2.0pre2 and duplicate reads were marked using MarkDuplicates in picard 2.23.8. Variant calls were generated via freebayes^53^ v1.3.4, keeping variants with a quality score above 100. Variants were annotated with gnomAD^54^ allele frequencies using SnpSift^54^ 5.0e. Known variants (i.e. those present in gnomAD) were retained for haplotype analysis.

### DMRs

DMRs were identified using Methylartist DSS with the HEK293T VCF file. DMRs were considered to be regions of at least 300bp with a >0.5 difference in aggregate methylation between alleles. Nanopore WGS reads were phased using WhatsHap^55^ phase and haplotag commands.

### CTCF peak calling

CTCF Nanopore-DamID peaks were called by first binning reads from both replicates separately into GATC fragment bins using BEDtools^56^ coverage. To remove bias from amplified regions of the HEK293T genome, we normalised the Nanopore DamID coverage by readcount and HEK293T Nanopore WGS coverage. The two replicates were then quantile normalised and averaged. Peaks were called on the average coverage using the DamID peak caller^47^ (https://github.com/owenjm/find_peaks). Peaks were only considered if they were present in both WGS-normalised and unnormalised peak sets.

### Code availability

CpG methylation analysis was performed using megalodon, available from https://github.com/nanoporetech/megalodon and Methylartist^37^, available from https://github.com/adamewing/methylartist.

## Data availability

Processed dataset are deposited in the Gene Expression Omnibus (GEO, GSE160383). Raw nanopore sequencing data and HEK293T Illumina WGS data are deposited in the NCBI Short Read Archive (SRA) repository as BioProject PRJNA850798.

## Competing interests

S.W.C’s registration, travel and accommodation at the 2022 Lorne Genome conference were paid for by Oxford Nanopore Technologies.

## Funding

This study was funded by an Australian Government Research Training Program (RTP) Scholarship to Y.M.A.J., the Australian Department of Health Medical Frontiers Future Fund (MRFF) (MRF1175457 to A.D.E.), the Australian National Health and Medical Research Council (NHMRC) (GNT1173711 to G.J.F., GNT1176574 to N.J. and GNT1161832 to S.W.C.), a CSL Centenary Fellowship to G.J.F., a UQ Genome Innovation Hub grant to S.W.C., and by the Mater Foundation (Equity Trustees / AE Hingeley Trust).

## Authors’ contributions

S.W.C. designed the study. S.W.C., Y.M.A.J, S.B.A., and M.K. performed the experiments. S.W.C., A.D.E., and G.J.F. performed the analysis. S.W.C. and G.J.F. funded the study. S.W.C., Y.M.A.J., N.J. and G.J.F. wrote the manuscript. All authors read and approved the final manuscript.

## Acknowledgements

The authors would thank C. James and A. Sehgal for technical assistance, and acknowledge the Translational Research Institute (TRI) for research space and equipment that enabled this research. We would particularly like to thank the TRI microscopy facility for assistance with this study. The authors thank the University of Queensland Genome Innovation Hub for continuing support.

---

**Supplementary figure 1:**
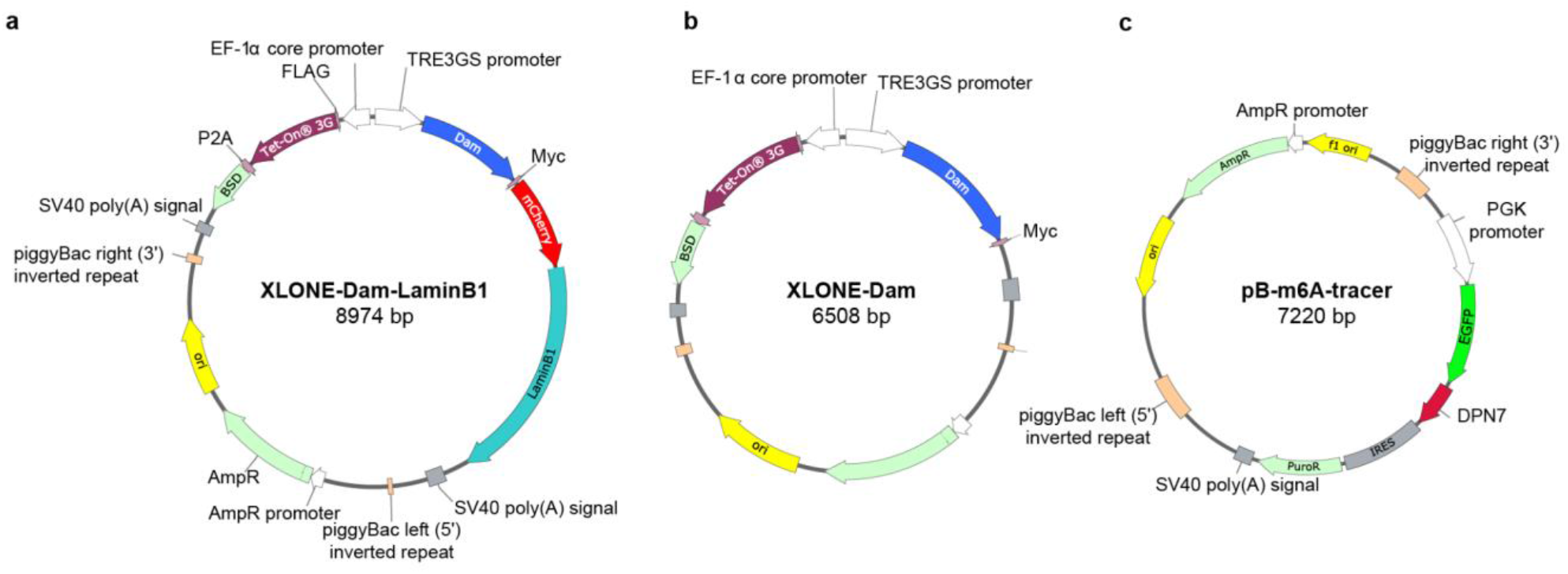
Constructs used for Nanopore-DamID. **a**, An all-in-one TetON-3G piggyBac construct based on XLONE^57^ enables generation of a stable cell line with an integrated doxycycline-inducible Dam-LaminB1. **b**, An all-in-one TetON-3G piggyBac construct based on XLONE^57^ enables generation of a stable cell line with an integrated doxycycline-inducible Dam. **c**, A piggyBac construct enables generation of a stable cell line constitutively expressing a catalytically inactivate DpnI fragment (Dpn7) fused to eGFP. Dpn7-eGFP binds to methylated GATC sites enabling visualisation of Dam-LaminB1 activity.

**Supplementary figure 2:**
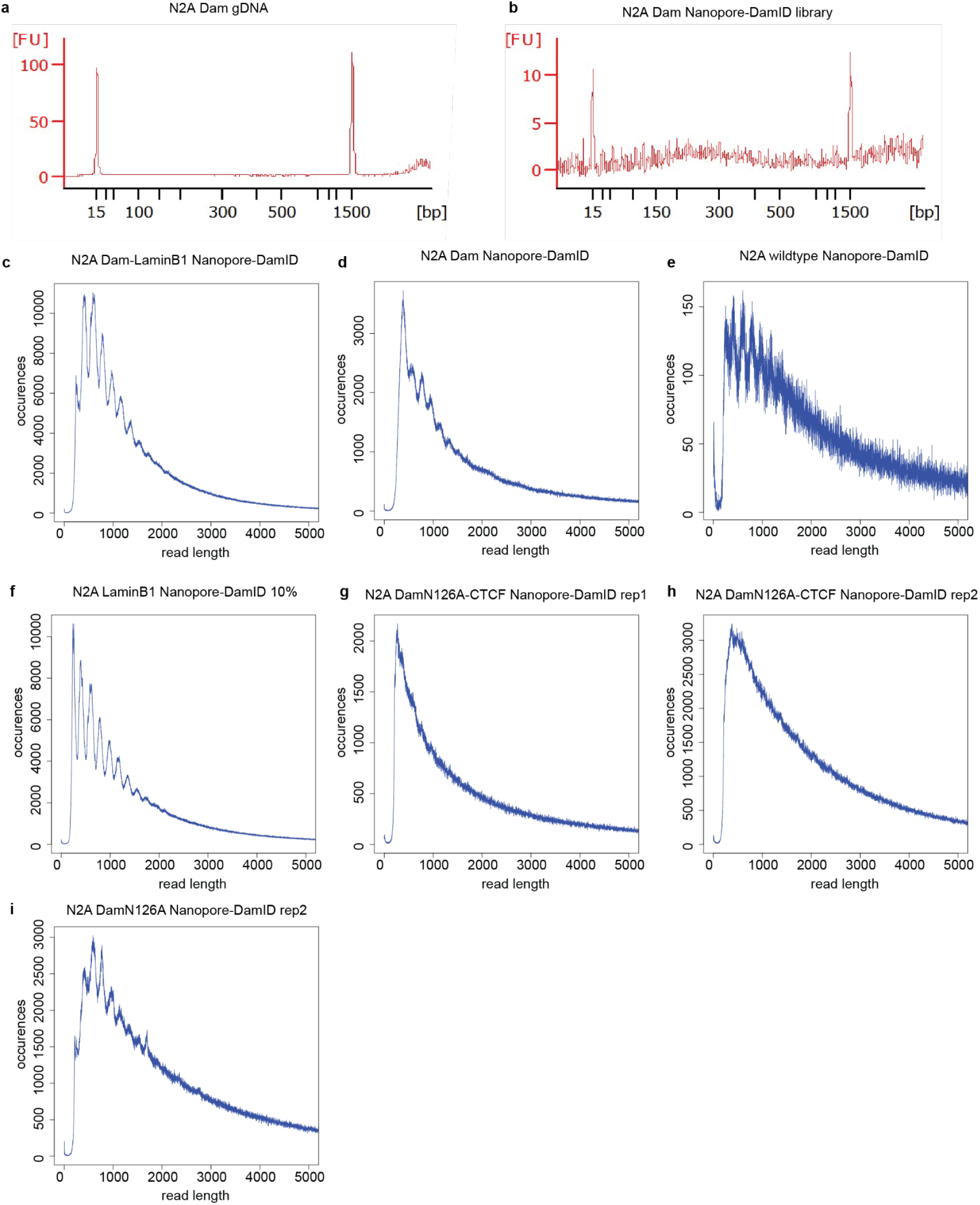
Size distribution of Nanopore-DamID libraries. **a**, An Agilent(R) tapestation trace of the size distribution of undigested gDNA from Dam-expressing N2A cells. **b**, A final Nanopore-DamID from Dam-expressing N2A cells. **c**, Fragment length distribution of reads from N2A LaminB1 Nanopore-DamID **d**, Fragment length distribution of reads from N2A Dam Nanopore-DamID. **e**, Fragment length distribution of reads from N2A wild type Nanopore-DamID. **f**, Fragment length distribution of reads from N2A LaminB1 Nanopore-DamID 10% in wild type cells. **g**, Fragment length distribution of reads from HEK293T CTCF Nanopore-DamID rep1. **h**, Fragment length distribution of reads from HEK293T CTCF Nanopore-DamID rep2. **i**, Fragment length distribution of reads from HEK293T DamN126A Nanopore-DamID.

**Supplementary figure 3:**
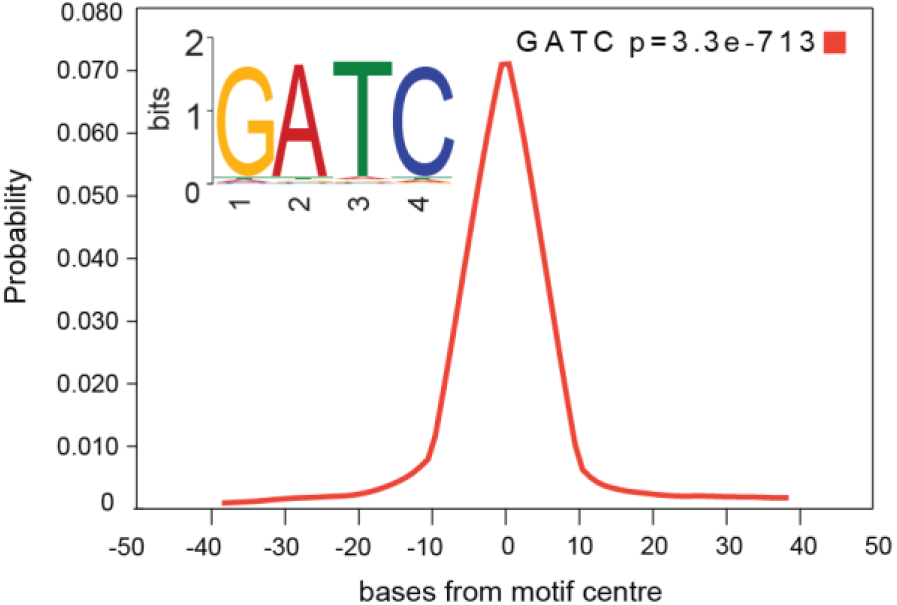
*De novo* discovery of GATC motif at the ends of Nanopore-DamID fragments. *De novo* motif discovery at the ends of Nanopore reads using MEME-ChIP^46^ revealed the motif GATC was strongly enriched.

**Supplementary figure 4:**
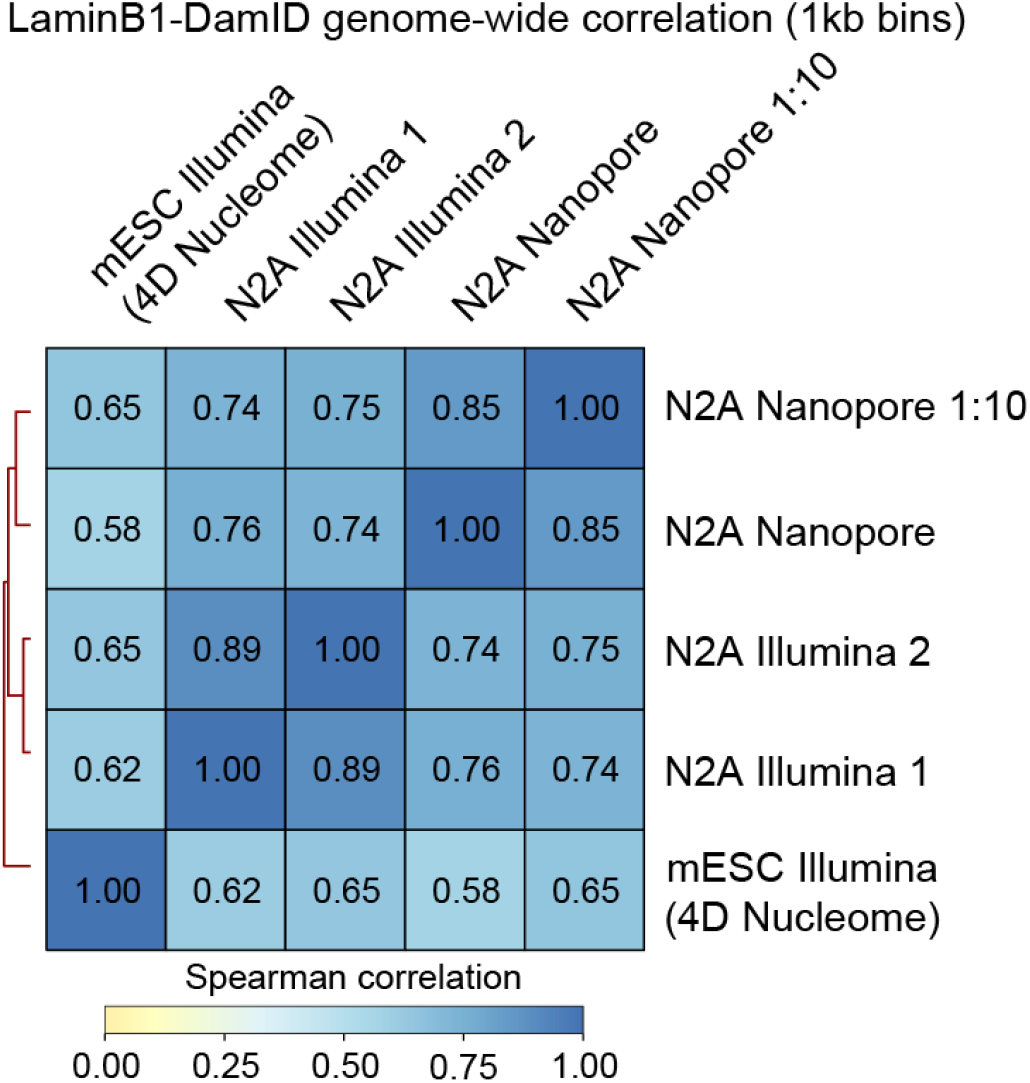
Correlation between Nanopore-DamID and conventional DamID datasets. Correlations were calculated genome-wide using deeptools^45^.

**Supplementary figure 5:**
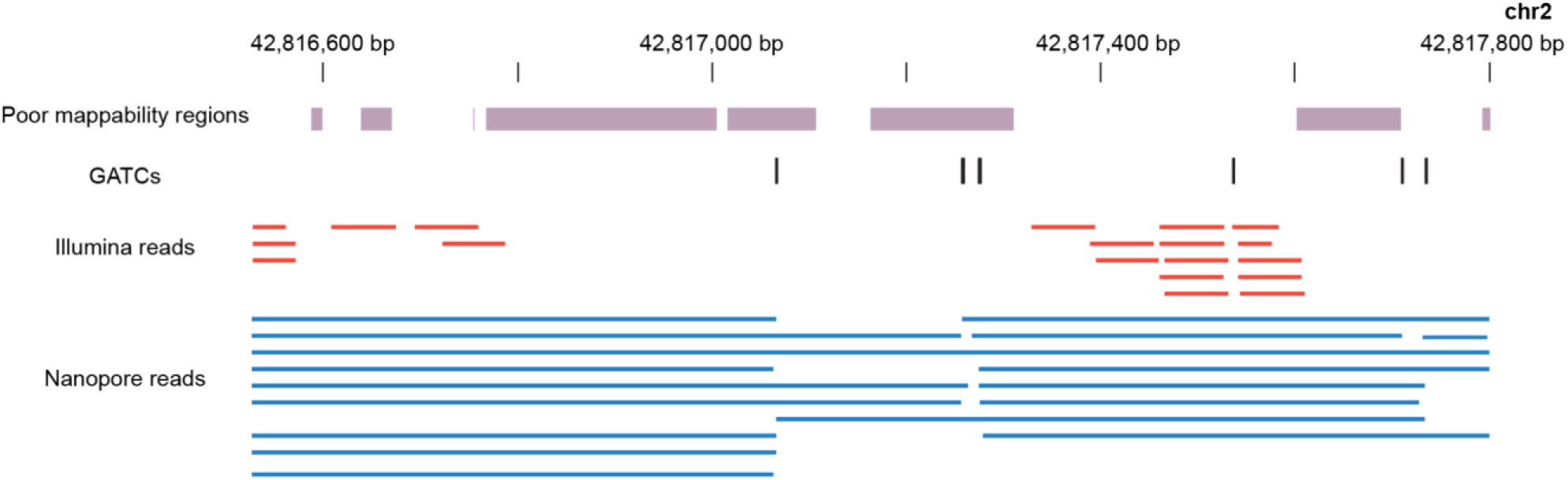
Superior alignment of Nanopore-DamID reads to regions of low mappability. Nanopore reads span regions that are unmappable for 75bp Illumina reads (identified by RSEG^58^).

**Supplementary figure 6:**
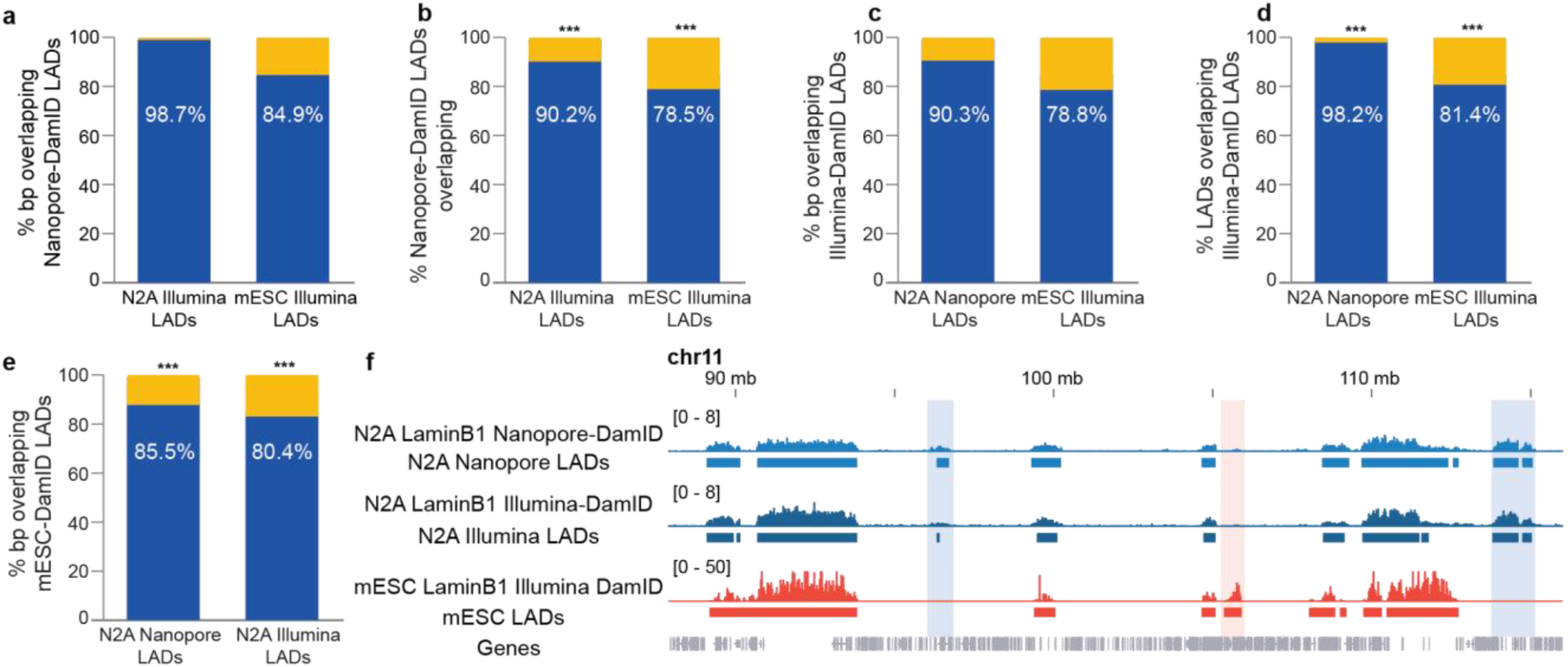
Comprehensive comparison of Nanopore and Illumina DamID. **a**, Proportion of bases of N2A-Illumina and mESC LADs that overlapped Nanopore-DamID consensus LADs. **b**, Proportion of N2A-Illumina and mESC LADs that overlapped Nanopore-DamID consensus LADs. **c**, Proportion of bases of N2A-Nanopore and mESC LADs that overlapped N2A Illumina-DamID consensus LADs. **d**, Proportion of N2A-Nanopore and mESC LADs that overlapped N2A Illumina-DamID consensus LADs. **e**, Proportion of bases of N2A Illumina and Nanopore LADs that overlapped mESC Illumina-DamID LADs. **f**, Some LADs were cell-type-specific LAD.

**Supplementary figure 7:**
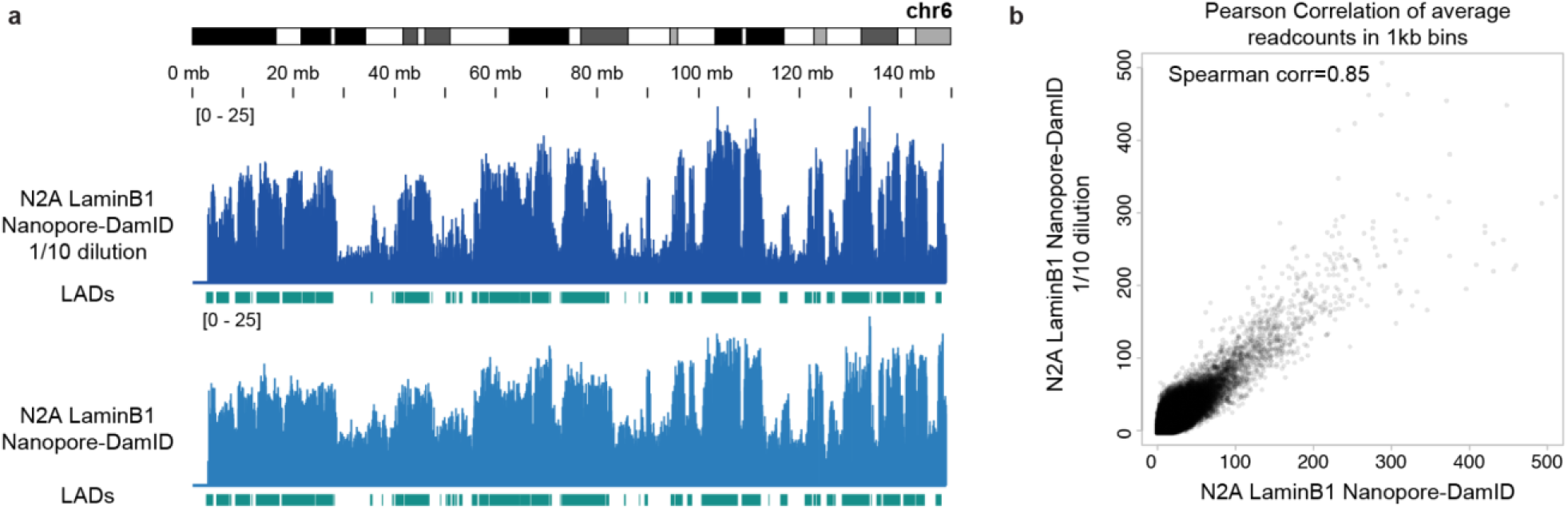
Selective sequencing of Lamin-associated DNA from a heterogeneous mixture of labelled and unlabelled cells. **a**, Comparison of LaminB1 Nanopore-DamID coverage from a 1/10 dilution of labelled cells in unlabelled cells on chromosome 6. **b**, Genome-wide correlation of diluted and undiluted Nanopore-DamID experiments.

**Supplementary figure 8:**
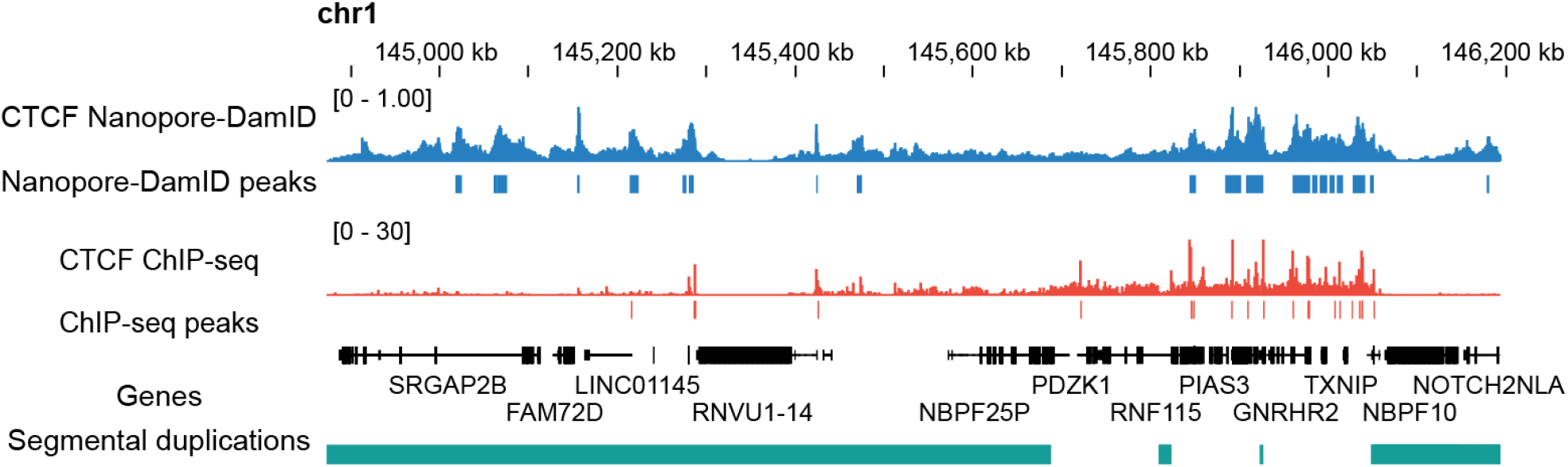
Nanopore-DamID detects occult CTCF binding sites in segmental duplications. CTCF Nanopore-DamID detected peaks in a segmental duplication on chromosome 1 that are not detected by CTCF ChIP-seq.

**Supplementary figure 9:**
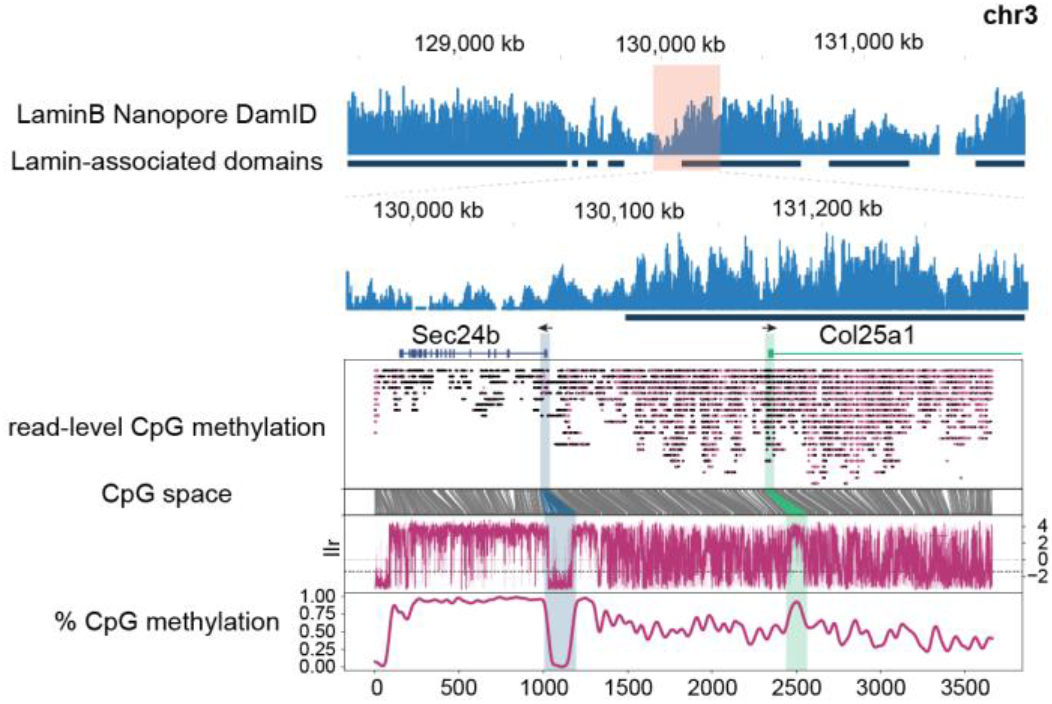
Resistance of a TSS on chr3 to LAD hypomethylation. An Example of the interaction of LaminB1 occupancy with DNA-methylation.

## Notes

### Competing Interest Statement

Registration, travel and accommodation at the 2022 Lorne Genome conference for S.W.C were paid for by Oxford Nanopore Technologies.

### Summary of Updates

The manuscript is updated with the inclusion of profiling of the transcription factor CTCF and further controls.

